# Offline tDCS modulates prefrontal-cortical-subcortical-cerebellar fear pathways in delayed fear extinction

**DOI:** 10.1101/2019.12.18.880658

**Authors:** Ana Ganho-Ávila, Raquel Guiomar, Daniela Valério, Óscar F. Gonçalves, Jorge Almeida

## Abstract

Transcranial direct current stimulation (tDCS) has been studied to enhance extinction-based treatments for anxiety disorders. However, the field shows conflicting results about the anxiolytic effect of tDCS and only a few studies have previously observed the extinction of consolidated memories.

Off-line tDCS modulates subsequent fear response (fear recall and fear extinction) neural activity and connectivity, throughout changes in the fear pathway that is critically involved in the pathogenesis of anxiety disorders.

Thirty-four women participated in a two-day fear conditioning procedure. On day 1, women were randomly assigned to the control group (n=18) or the tDCS group (n=16) and went through a fear acquisition procedure. On day 2, the tDCS group received 20min tDCS at 1mA [cathode – F4; anode – contralateral deltoid] immediately before extinction and while inside the MRI scanner. The control group completed the extinction procedure only.

fMRI whole brain contrast analysis showed stimulation dependent activity patterns with the tDCS group showing decreased neural activity during the processing of the CS+ and increased activity during the processing of the CS, in prefrontal, postcentral and paracentral regions, during late extinction. PPI analysis showed tDCS impact on the connectivity between the left dorsolateral prefrontal cortex and three clusters along the cortical–amygdalo–hippocampal– cerebellar pathway, during the processing of the CS+ in late extinction (TFCE corrected at p <.05).

The increased neuronal activity during the processing of safety cues and the stronger coupling during the processing of threat cues might well be the mechanisms by which tDCS contributes to stimuli discrimination.

**Highlights:**

- The anxiolytic effect of cathodal tDCS is controversial.
- We show cathodal tDCS modulatory effect on delayed extinction of the fear response.
- Cathodal tDCS modulates the processing of safe and threatening cues.
- Cathodal tDCS modulates the activity and connectivity of the fear network.

## 1. Introduction

Anxiety disorders represent the most prevalent psychiatric conditions worldwide (Baxter et al., 2013; Craske et al., 2017), negatively impacting individual lives, and impairing functioning in professional and social contexts (Craske and Stein, 2016). Notwithstanding the diversity and accessibility of pharmacological treatments targeting anxiety disorders, those currently available show limited efficacy (Baldwin et al., 2011) and present constraints with respect to tolerability and safety profiles (Arcoraci and Spina, 2015). As for psychological treatments, despite their increased tolerability, their efficacy rates are no better than pharma (Carpenter et al., 2018; Loerinc et al., 2015) and are time-consuming. Hence, albeit the combination of medication with psychological treatments seems to be an option, it still presents limited therapeutic gains (e.g. (Keller et al., 1994; MB et al., 2000)). Aiming at exploring transcranial direct current stimulation (tDCS) effect as an add-on medication-free treatment to decrease anxiety related symptoms, in this translational study we mapped its impact during experimental fear extinction.

Non-invasive brain stimulation (NIBS) strategies such as tDCS have shown promising effects in decreasing different psychiatric symptoms, either as a stand-alone or combined with pharmacotherapy and psychological interventions (Donse et al., 2018). Among NIBS, tDCS is the most portable and user-friendly technique, displaying a high safety and comfort profile (Nikolin et al., 2018). tDCS consists of a direct current delivered through electrodes aimed at changing cortical excitability by increasing or decreasing neuronal membrane polarity (Nitsche and Paulus, 2000) or by altering the oscillatory activity of a particular network (Ikeda et al., 2019).

The effectiveness of tDCS in managing symptoms of anxiety has been anecdotally suggested in pre-clinical and clinical studies (Carnevali et al., 2019; Pedro Shiozawa et al., 2014a, 2014b). More extensively, several laboratorial studies have used tDCS to study classical conditioning of the fear response - the translational model for the development, maintenance and elimination of anxiety symptoms. According to this model, the fear response is elicited by environmental threats through an associative learning whereby the neutral stimulus (conditioned stimulus, CS+) is paired with an aversive stimulus (unconditioned stimulus, US) and a danger predicting value is assigned to it, eliciting the conditioned response (Rachman, 1977). According to the model, a deregulated fear response may lead to incorrect detection of danger and sustained anxiety-related symptoms (e.g. hyper-vigilance and excessive worrying). Thus, to eliminate the consolidated CR, extinction (the translational equivalent to exposure-based therapies) is implemented, consisting of consecutive presentations of the CS+ alone, leading to a new learning that in turn inhibits the CS+/US association (Lonsdorf et al., 2019).

By delivering tDCS to the right or left DLPFC, researchers aimed to enhance fear extinction by artificially modulating brain activity to decrease the fear response (Asthana et al., 2013; Dittert et al., 2018; Ganho-Ávila et al., 2019a; Mungee et al., 2016). Counterintuitively, whereas previous studies have shown that tDCS using the anode electrode over the DLPFC leads to decreased fear, this gain was also associated with the adverse effect of decreasing stimuli discrimination (Dittert et al., 2018). Differently, using the cathodal electrode over the DLPFC led to no effect in decreasing fear but seemed to result in increased stimuli discrimination (Abend et al., 2016; Ganho-Ávila et al., 2019a). Such perplexing findings motivate our study, by suggesting the need to better understand the mechanism by which cathodal tDCS interferes with the fear network.

tDCS over the right DLPFC-contralateral deltoid montage was used in previous case studies showing tDCS effectiveness in reducing anxiety related symptoms (Pedro Shiozawa et al., 2014a, 2014b). This finding is understandable as the DLPFC is known to be the hub region associated with the appraisal of fear experiences (Ertl et al., 2013) besides being involved in fear extinction learning (Fullana et al., 2018). In particular, the rDLPFC is interconnected with the ventro medial prefrontal cortex (vmPFC) to downregulate the amygdala activity through emotion regulation and value detection (Phelps et al., 2004). Moreover, current modelling literature shows that increased current intensity does not occur under the electrode but rather between electrodes (Woods et al., 2016), allowing for the modulation of the entire pathway that relies between electrodes and beyond, interfering not only with cortical structures but also with subcortical structures.

During fear extinction, participating regions compose the prefrontal cortical-amygdalo-hippocampal-cerebellar pathway. Previous meta-analysis (Fullana et al., 2018) show that the rostro-dorsal cingulate cortex, the medial prefrontal cortex, the bilateral anterior insula (extending to the frontal operculum), the dorsolateral PFC, the basal ganglia, the anterior and medial thalamus and the periaqueductal grey area are consistently activated during extinction learning. However, when looking at late extinction in particular, the anterior thalamus, the ventral putamen and the right anterior insula prevail. Further, when fear extinction occurs in a distinctive context from fear acquisition the bilateral middle occipital cortex and the left somatosensory-supramarginal cortices are consistently engaged during extinction. This contrasts with the pattern of neural activation when extinction occurs in the same context as acquisition where the left anterior and the right posterior insula are engaged along with the anterior cerebellum. In fact, previous fMRI studies have also suggested the cerebellum participation in the extended fear network, associated with the regulation of the autonomic response during both early (Fullana et al., 2018) and late phases (Kattoor et al., 2013) of fear extinction, and in the processing of time, valence and prediction of stimuli contingency (for in-depth literature we recommend the systematic review by Lange and colleagues (Lange et al., 2015)).

Although the hippocampus is broadly known as responsible for processing context features during the acquisition of the fear response (Alvarez et al., 2008; Marschner et al., 2008), a recent face-to-face meta-analysis between studies conducted in the same vs distinct contexts during extinction, did not show a consistent involvement of this region (Fullana et al., 2018).

Besides fMRI studies, electrophysiological studies have also shown the fear pathway extension. For example, the artificial induction of synchronized theta frequencies lead by the prefrontal cortex produces a coherent corresponding spike firing in the hippocampus and the amygdala (Lesting et al., 2013, 2011). These coherent theta firing are associated with processes of appraisal and experience of fear (Ertl et al., 2013), fear extinction learning (Lesting et al., 2013, 2011) or fear extinction reconsolidation (Narayanan et al., 2007). Also, the reduction of gamma power in the ACC and in the vmPFC is correlated with fear extinction (Mueller et al., 2014).

The translational value of extinction for anxiety disorders relies on the specificities of delayed extinction, which allows for the temporal gap between learnings (the fear response acquisition and the fear response extinction), offering the needed time window for the consolidation mechanisms associated to memory and learning processes (Schwarzmeier et al., 2019) and mimicking real-life symptom onset and treatment. However, most previous studies have used within-session acquisition and extinction learning procedures not allowing for a consolidation period. Consequently, what has been claimed the laboratory analogue for exposure-based therapies has, in fact, targeted recent memories which are known to present distinctive neural patterns and to be qualitatively more susceptible to interference (McClelland, 2013).

Because long-term consolidated memories are a typical feature of treatments in anxiety disorders, to understand how tDCS interacts with the network when processing delayed extinction is of the utmost importance; otherwise, it compromises translational findings. We aimed to fill this gap by implementing a two-day fear conditioning procedure and observing how a single 20-min tDCS session over the rDLPFC interferes with the neural activity and connectivity patterns of the prefrontal cortical–amygdalo–hippocampal–cerebellar pathway during delayed extinction. We hypothesized that off-line tDCS over the rDLPFC would lead to activity and connectivity changes in the entire pathway critically involved in the pathogenesis of anxiety disorders, from the frontal regions up to the cerebellum.

## 2. Materials and Methods

### 2.1 Participants

Forty-eight women were recruited to take part in the experiment, having given their informed consent according to the local ethical committee and compliant with the Declaration of Helsinki recommendations. Three participants withdrew from the study after feeling discomfort during the experimental task. Data from eight participants were excluded because they did not acquire the fear response, and data from three participants were excluded due to technical problems. As such, 34 women (mean age of 23.32 years, SD = 5.67, min = 18, max = 38) completed the experiment, and were assigned to either the control (N=18), or tDCS group (N = 16). Data from one participant concerning US intensity, and ASI-3-PT, and from two participants in BSI were not recorded due to technical problems in Eprime (remaining data from these participants were still included in the analysis); for details see Table S1in supplementary materials). In day 1, participants showed high but below clinical significance mean scores in what concerns anxiety state, anxiety trait and anxiety sensitivity self-report measures. No differences were found between the two groups 1 (*p* = .71, *p* = .60, and *p* = .28, respectively)

Exclusion criteria for participation included: (1) history of psychiatric disorders; (2) use of psychoactive medication; (3) pregnancy; (4) caffeine and/or alcohol intake 24 hours before sessions; (5) physical exercise two hours before sessions; (6) having had a meal two hours before sessions (Boucsein et al., 2012); (7) auditory or visual (non-corrected) deficits; and (8) the typical exclusion criteria for MRI scanners. Additionally, participants had to show fear acquisition on Day 1, to allow for observations on fear extinction processes on Day 2. Based on previous literature, the sample size was established at a minimum of 15 subjects per group (Agren, T., Engman, J., Frick, A., Bjorkstrand, J., Larsson, E.-M., Furmark, T., & Fredrikson, 2012; LaBar et al., 1998; Phelps et al., 2004).

### 2.2 Procedure

Participants completed a partial-reinforcement auditory fear conditioning procedure for two consecutive days, allowing for the fear memory to consolidate (day 1 – psychological assessment, habituation to stimuli, fear response acquisition; day 2 – [tDCS session for the tDCS group, followed by] fear extinction). Before starting, participants underwent a brief assessment to cover for the exclusion criteria. Also, the volume intensity of the US was individually set. The fear acquisition procedure occurred in the laboratory and the delayed fear extinction procedure inside the MRI scanner. For the first day, we collected skin conductance responses (SCRs) to measure the autonomic fear response, as well as self-report ratings on valence, arousal contingency and expectancy. For the second day we collected structural and functional MRI data.

#### 2.2.1 Stimulus

Three stimuli were presented – the unconditioned stimulus (US), the conditioned stimulus (CS+), and the control stimulus (CS-; never paired with the US). As for the US, we used a woman’s scream selected from the International Affective Digitized Sound System (IAPS, item 277; (Bradley and Lang, 1999)). The US volume was individually set in the first session between 90-96dB (according to the participants’ comfort level), testing step-wise with a dummy aversive sound (IAPS, item 276; (Bradley and Lang, 1999)) added by a Visual Analog Scale for pain to measure discomfort (Huskisson, 1974) while the sound was delivered through noise-cancelling headphones. In the session proper, the US was presented for 2s. The CS+ and CS-were 12×12cm squares either blue or yellow (colours were counterbalanced across participants), with a white background, presented in a DELL P2012H monitor for 4s across sessions. Across sessions, we pseudo-randomized the CS+ and CS-trials such that no more than two consecutive presentations of the same category were allowed.

We used E-Prime (2.0.10.353 Standard SP1, Psychology Software Tools, Pittsburgh, PA) scripts to design, collect and analyse data. US triggers and onset markers were sent to a second computer where we collected the electrodermal activity across habituation and acquisition.

#### 2.2.2 Day 1 – Habituation and fear acquisition

Participants were randomly assigned to the experimental groups (tDCS or control). The habituation phase consisted of 8 non-reinforced CS+ and 8 CS-presentations. The acquisition phase consisted of a partial reinforcement procedure at 75% (i.e. 12 out of 16 CS+ presentations were paired with the aversive sound). US presentations overlapped with the last 2s of the CS+. In total, there were 16 CS+ and 16 CS-trials, each followed by a jittered interstimulus interval (ISI; a black fixation cross presented in the centre of the screen for 10 to 12s). The acquisition of the fear response was considered successful whenever in the late phase of acquisition (last 5 trials) the mean SCRs to the CS+ was higher than the mean SCRs to the CS-. For those showing earlier habituation to the US in the late phase, we alternatively considered the middle phase (from the 6^th^ to 10^th^ trial inclusively). We assumed .01 μS as the minimum difference (Oyarzún et al., 2012). Participants that showed fear response on day 1 were invited for the second day and were instructed not to use hair products or have wet hair for the next session to avoid interference with tDCS stimulation.

#### 2.2.3 Day 2 – tDCS session and delayed fear extinction procedure

Participants assigned to the tDCS group started day 1 by completing the 20-minute tDCS session (including 30s of ramp up and 30s of ramp down), with a current intensity of 1mA, delivered through a tDCS 1-channel stimulator (TCT Research Limited, Hong Kong). The cathode was placed over the right DLPFC (F4, 10/20 international system) and the anode over the contralateral deltoid muscle (extra-cephalic montage). The sponges covering the electrodes were 5×5cm in size and were saturated with 10 mL of 0.9% saline solution immediately before the stimulation session. During the offline stimulation session, the participants were instructed to stay still and calm and wait until the end of the session. Offline tDCS was preferred over online tDCS, due to previously reported MRI artefacts caused by the presence electrical current inside the MRI scanner (Antal et al., 2014). tDCS adverse effects were assessed immediately following stimulation (Table S1 in supplementary materials) and participants were then led to the extinction procedure inside the MRI scanner. Participants assigned to the control group went directly to the extinction session inside the MRI scanner. Before starting extinction, to test fear acquisition, participants were asked to recall the colour of the square paired with the US on the acquisition session. This procedure additionally builds up on fear recall procedures to boost reconsolidation (Schiller et al., 2018, 2010), and no further information was offered about the auditory stimuli. Inside the MRI scanner all participants wore headphones to simulate the experimental setting of day 1 and allow for an equal expectation of hearing the US during the extinction session. The extinction session consisted of 10 CS+ and 10 CS-trials. Each trial consisted of a fixation cross presentation for 16s followed by the CS+ or CS-presentation for 4s. During extinction, the US was never presented. The stimuli were presented using an Avotec projector, controlled by “A Simple Framework” (Schwarzbach, 2011) under MATLAB R2014a (The MathWorks Inc., Natick, MA, USA). The presentation was then back presented through a mirror attached to the head coil. Participants viewed the images passively in a pseudo-random order (no more than two consecutive presentations of the same colour in a row) for 7 minutes.

### 2.3 Data Acquisition, pre-processing and analysis

#### 2.3.1 Psychological questionnaires

Data on baseline psychological status according to the Anxiety Sensitivity Scale −3-PT (Ganho-Ávila et al., 2019b), the Behavioural Symptoms Inventory (Canavarro, 2007), the State Anxiety Inventory (STAI-1) and Trait Anxiety Inventory (STAI-2; (Spielberger, 1989) were collected using the experiment computer, screen and keyboard.

#### 2.3.2 Skin conductance responses

We collected electrodermal activity on Day 1, using a pair of Powerlab 26T finger electrodes (MLT116F; ADInstruments, Ltd., Dunedin, New Zealand) attached to the medial phalanges of the index and middle fingers of the left hand. These electrodes were connected to a galvanic skin signal Amplifier (FE116; ADInstruments, Ltd., Dunedin, New Zealand) that collected data every 200 ms, filtering out frequencies above 50 Hz. The signal was pre-processed in MATLAB (2013, The MathWorks, Inc, Natick, Massachusetts, United States) using in-house scripts. To use all trials and not confound with US responses (starting at 2s after the CSs onset in 75% of the trials), we calculated the SCRs on a trial-by-trial basis using the first 2s after the onset of each stimulus. The 500 ms pre stimulus onset were used as baseline. For the SCRs, the first peak minus baseline values per trial were square rooted (Oyarzún et al., 2012). Responses starting before stimulus presentation were considered artefacts and were manually deleted. The first presentation of each stimulus was deleted to account for orienting response, so 7 trials in habituation and 15 in acquisition and extinction phases will be reported.

#### 2.3.3 Self-report measures

At the end of habituation and acquisition sessions, we collected offline valence and arousal self-reports per stimuli using Lang’s (1980) nine-point Self-Assessment Manikin (SAM) scales (Lang and Bradley, 2010). For valence, we used a 1-9 scale (1 = *highly unpleasant*; 9 = *highly pleasant*); for arousal we used a 1-9 scale (1 = *highly calm*; 9 = *highly excited),* for the expectancy of US presentations paired with the CSs presentation we used a 0-9 scale (0 = *I was sure the sound was not coming*; 9 = *I was sure the sound was coming*; finally for the contingency awareness concerning the CSs/US association we used a 0%-100% scale, in increments of 25% (0% = *the CS was never paired with the US*,; 100% = *the CS was always paired with the US* (Ganho-Ávila et al., 2019a). Participants were asked to register their answers with the use of the keyboard.

#### 2.3.4 MRI data

MRI data was collected on a 3T Siemens Magnetom Trio Tim scanner with a 12-channel head coil. We collected high-resolution structural T1 weighted images, using a MPRAGE pulse sequence (repetition time [TR] = 2530 ms; echo time [TE] = 3.29 ms; flip angle: 7 degrees; field of view [FOV] = 256 × 256 mm; matrix size: 256 × 256; voxel size: 1×1×1 mm; slice thickness = 1 mm; number of slices = 192 ascending interleaved). An EPI pulse sequence was used for T2* contrast (TR = 2000 ms; TE = 30 ms; flip angle: 90°; FOV = 256 × 256 mm; matrix size: 64×64; voxel size = 4×4×4 mm; slice thickness = 4 mm; number of slices = 30 ascending interleaved even-odd slices). We discarded the first two volumes of each run allowing for signal equilibration. For pre-processing proposes, we used SPM12 software version 7219 and in-house MATLAB scripts.

#### 2.3.4.1 Definition of regions of interest (ROIs)

To understand the activity and connectivity of the fear network, and how this is modulated by tDCS, we defined a set of cortical regions of interest. Considering that tDCS was delivered over the DLPFC, that is interconnected with the vmPFC which in turn is engaged in mechanisms of downregulation of the amygdala (Fullana et al., 2018), the bilateral DLPFC, and the vmPFC were selected using neurosynth.org data (http://neurosynth.org/). Bilateral DLPFC were defined from automated metaanalysis out of 362 records, using the term “DLPFC” and the reverse inference method. We created a spherical ROI of 8mm radius using SPM toolbox Marsbar (http://marsbar.sourceforge.net/), centred over the significant cluster (Montreal Neurological Institute [MNI] coordinates; xyz = + - 42, 34, 32) and checked its overlap with our stimulation site (F4) using *tool mni2tal* (Yale Bioimage suite; http://sprout022.sprout.yale.edu/mni2tal/mni2tal.html). To define the vmPFC we followed the same pipeline, using the term “vmPFC” and the reverse inference method. The vmPFC was defined from automated meta-analysis out of 199 records, creating a spherical ROI with 8 mm radius centred over the significant cluster (MNI coordinates; xyz= 0, 30, −16).

A grey matter mask was used to reduce the number of multiple comparisons in each analysis (MNI grey matter mask template downloaded from Brain Imaging Centre, Montreal Neurological Institute, McGill University - https://digital.lib.washington.edu/researchworks/handle/1773/33312).

All masks were normalized into MNI space, binarized using SPM12 and finally checked for correct alignment.

#### 2.3.4.2 Functional magnetic resonance imaging

SPM12 was used to analyse fMRI data (version 7219; https://www.fil.ion.ucl.ac.uk/spm/) in Matlab environment (9.3.0.713579 R2017b; https://www.mathworks.com/). Before pre-processing, we manually set the coordinates of the anterior commissure of the anatomical scan to zero and applied the same transformations to the functional scans of every subject. The pre-processing of the functional data included slice-timing correction (sync interpolation), motion correction with respect to the first volume of the first functional run with a 7th degree B-spline interpolation, coregistration of the anatomical scan to the first volume of the first functional run on an individual basis. Functional data were smoothed using a Gaussian kernel (FWHM of 8 mm) and interpolated to 3 × 3 × 3 mm voxels.

The general linear model (GLM) was used to fit beta estimates to the events of interest. The default high-pass filter cut-off of 128 seconds was included. The first derivatives of the six motion parameters describing volume-to-volume motion were added (not convolved) as predictors of no interest to attract variance attributable to head movement. Experimental events were convolved with a canonical hemodynamic response function.

Based in the literature (Lonsdorf et al., 2019, 2017) we defined four regressors of interest: CS^+^early, CS^+^ late, CS^-^early and CS^-^late, where the early phase included the first five and the late phase the last five trials. Model parameters were estimated using the classical Restricted Maximum Likelihood (REML) method, with an autoregressive AR(1) to account for serial correlations. Group-level significance testing was performed using Monte Carlo simulations (with the CoSMo Multivariate Pattern Analysis toolbox [CoSMoMVPA] in MATLAB) with 10,000 iterations, corrected for multiple comparisons using Threshold-Free Cluster Enhancement (TFCE; (Smith and Nichols, 2009).

##### Whole brain contrast analysis

To understand how the neural activity during fear extinction is modeled by tDCS, we performed whole brain between group contrast analysis. Phase-dependent contrasts (CS+ early *vs.* CS+ late; CS-early *vs*. CS-late) and condition-dependent contrasts (CS+ early *vs*. CS-early; CS+ late *vs*. CS-late; early phase = first five trials; late phase = last five trials) were performed. The *a priori* hypothesis defined that there would be distinctive condition-dependent (CS+ *vs*. CS-; increased or decreased stimuli discrimination) and phase-dependent (early *vs*. late) neural activity between groups (tDCS and control).

##### Contrast psychophysiological interaction (PPI) analysis

To understand the CSs-specific connectivity patterns between the cortical ROIs (DLPFC, and the vmPFC) and the whole brain grey matter mask during the early *vs*. late phase of extinction, we estimated the neural activity differences of time correlations between conditions using contrast PPI analysis (Di et al., 2018; Friston et al., 1997; Gitelman et al., 2003; O’Reilly et al., 2012) in SPM12. We extracted the first eigenvariate timeseries data per seed ROI, controlling for all conditions of interest (F contrast). The resulting deconvolved seed ROI time course was multiplied by the task time course (convolved with the haemodynamic response function) to form the PPI predictor (Di et al., 2018). We then computed one GLM with three regressors: the interaction term, the seed region’s extracted signal and the experimental condition (PPI.ppi, PPI.Y and PPI.P, respectively). After model estimation, a contrast map was created with the PPI interaction as the regressor of interest. Between-groups analyses (two sample *t-tests*; two-tailed) were performed with the toolbox CoSMoMVPA. The resulting group level statistical Z maps were thresholded at |z| > 1.96 (*p* < 0.05, TFCE corrected) and between groups results were Bonferroni corrected for multiple comparisons. Clusters were defined as significant contiguous groups of voxels regardless of its extension. The *a priori* hypothesis that off-line tDCS over the rDLPFC interferes with the fear network during delayed extinction, defined that distinctive connectivity patterns would be found in the prefrontal cortical-amygdalo-hippocampal-cerebellar pathways during extinction for the tDCS group when compared with the control group. No direction was predicted.

To further understand the association between day 1 self-report measures of arousal, valence, contingency and expectancy, and SCRs with day 2 neural activity, correlational analysis were conducted.

## 3. Results

We sought to understand how tDCS modulates the fear pathway during the extinction of a consolidated fear response. For such, we implemented a 2-day experimental procedure. On day 1, participants were split in two groups (the tDCS and the control group) and learned to fear the conditioned stimulus in the laboratory. To confirm fear acquisition, SCRs and self-reported arousal, valence, contingency and expectancy were collected. On day 2, participants went through extinction inside the MRI scanner, where the neural activity and connectivity of the fear network was observed.

### 3.1 Psychophysiological data

SCRs collected during the habituation and acquisition phases from day 1 confirmed the successful acquisition of the fear response. As expected, we found no differences in psychophysiological arousal between the CS+ and the CS-during the habituation phase, both for the control group (*t* (17) = −.38, *p* = .710, 95% IC [-.084, .058] and for the tDCS group (*t* (14) = −1.35, *p* = .198, 95% IC [-. 116, .026], suggesting that the CSs were equated at baseline in terms of the arousal elicited. After fear acquisition, the CS+ elicited more increased responses than the CS-, both for the control group (*t* (17) = 2.84, *p* = .011, 95% IC [.030, .205]) and for the tDCS group (*t* (14) = 2.54, *p* = .023, 95% IC [.011, .130]; (Figure 1).

**Figure 1.**
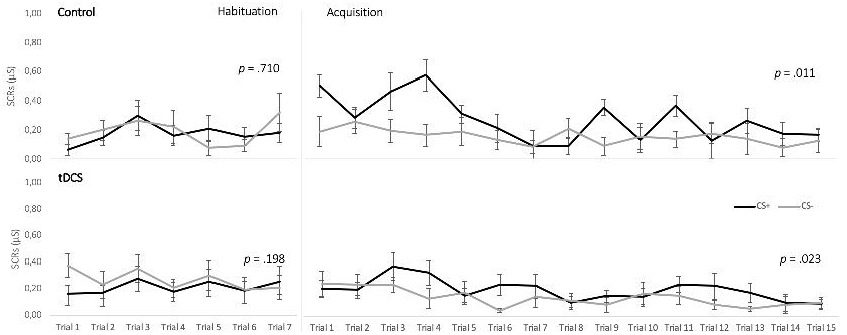
Day 1 SCRs. Day 1 SCRs for the habituation and acquisition procedures for the CS+ and the CS-per trial. CS+: conditioned stimuli; CS-: non-reinforced or control stimuli; tDCS: transcranial Direct Current Stimulation group; Control: control group. Error bars represent standard errors of the mean (SEM).

### 3.2 Behavioural data

Self-ratings data on valence and arousal of the CS+ and the CS-validate the psychophysiological data. After habituation, results showed no significant differences between the CSs for the control group for arousal (*t* (17) = .13, *p* = .897, 95% IC [-.838, .950]) and valence ratings: *t* (16) = −.112,*p* = .912, 95% IC [-1.172, 1.053]) and the same was true for the tDCS group (arousal: *t* (15) = .37,*p* = .718, 95% IC [-.600, .850]; valence: *t* (15) =-1.14,*p* = 2.71,95% IC [-1.433, .433]).

After acquisition, both groups evaluated the CS+ as evoking more arousal than the CS-(control group: *t* (16) = 7.90,*p* < .000, 95% CI [2.625, 4.551]; tDCS group: *t* (15) = 5.37,*p* < .000, 95% CI [1.923, 4.452]). Similarly, both groups evaluated the CS+ as being more negative than the CS-(control group: *t* (15) = −4.28,*p* = .001, 95% CI [-4.680, −1.570]; tDCS group: *t* (15) = −4.99,*p* < .001, 95% CI [-4.995, −2.005]).

Confirming that participants learned the association between the CS+ and the US, self-reports on CS+/US and CS-/US contingency after acquisition showed significant differences between stimuli, both in the control group (*t* (16) = 5.01,*p* < .000, 95% CI [1.42, 3.52]) and in the tDCS group (*t* (15) = 13.76,*p* < .000, 95% CI [2.48, 3.39]). For better visualization, Figure S1 in supplementary materials depicts self-reports’ data concerning post-habituation and post-acquisition for each group.

### 3.3 fMRI results

Whole brain contrast analysis showed two clusters where activity patterns were distinctive between groups during late extinction for the contrast CS+ > CS- . Cluster 1 (peak voxel MNI coordinates: −18, 15, 45; mean z = 2.07) included, among other areas, the frontal middle and the frontal superior left cortices. Cluster 2 (peak voxel MNI coordinates: −9, −33, 57; mean z = 2.00) included the left and right paracentral region and the right post central area (TFCE corrected at p < .05). Distinctive brain responses to the CSs per group are depicted in Figure 2. Whereas the tDCS group showed increased activity in both Cluster 1 and Cluster 2 during the processing of the CS-, the control group showed a decreased activity in Cluster 1 both during the processing of the CS+ and the CS-, and an increased activity during the processing of the CS+ only in Cluster 2 (for detailed information regarding the clusters’ activity patterns per region according to Automated Anatomical Labeling [AAL], see supplementary materials Table S2). Cluster 1 and Cluster 2 did not survive Bonferroni correction for multiple comparisons (p < .0125; Z > 2.241).

**Figure 2.**
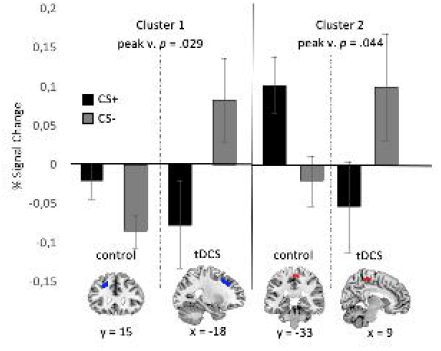
Mean Percentage signal change during late extinction per stimuli in control and tDCS group for cluster 1 (left; blue) and cluster 2 (right; red). Voxel-wise TFCE corrected at p > .05. Uncorrected for multiple comparisons when accounting for 4 contrasts. Control: control group; tDCS: transcranial Direct Current Stimulation group. peak v. *p: p*-value in the voxel with the maximum activity. CS+: conditioned stimuli; CS-: non-reinforced or control stimuli. Error bars represent standard error of the mean (SEM). Images displayed according to the neurological convention (left-right).

PPI analysis per seed ROI for the whole brain grey matter (Di et al., 2018; Friston et al., 1997), by behavioural task contrast (contrast 1: CS+early > CS+late, contrast 2: CS+early > CS-early, contrast 3: CS+late > CS-late, contrast 4: CS-early > CS-late) was used to inspect phase-dependent (early *vs*. late) and condition-dependent (CS+ *vs*. CS-) functional connectivity patterns of the fear network. The seed ROIs were the lDLPFC, the vmPFC and the rDLPFC (the stimulation site, according to our original hypothesis). To observe the impact of tDCS, we estimated two-sample t-tests for each analysis.

The contrast CS+late > CS-late was the single one showing significant differences, and these were found specifically in the connectivity between the lDLPFC and three clusters. Cluster 1 peak voxel was located in the right frontal superior region (7055 voxels; xyz=24, 54, 9, mean z = −2.89), Cluster 2 in the left temporal middle region (38 voxels, xyz=-44, 44, −24; mean z = −2.34), and Cluster 3 in the left middle frontal region (88 voxels; xyz = −30, 30, 36; mean z = −2.34; TFCE corrected at p < .05 voxel-wise and Bonferroni corrected accounting for four comparisons). Figure 3 shows the location and extension of each of the three clusters.

**Figure 3.**
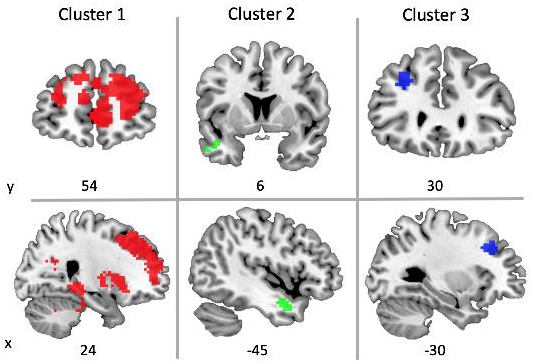
Two-sample t-tests for group differences on lDLPFC-whole brain PPI analysis, during the processing of the CS+ in late extinction. Cluster 1 (red), cluster 2 (green), cluster 3 (blue), were statistically significant at *p* < .05 (Bonferroni corrected). Top row: coronal view; bottom row: sagittal view; x, y = MNI coordinates. Images displayed according to the neurological convention (left-right).

Cluster 1 involved a broad set of brain regions known to have distinctive roles in fear extinction. To better distinguish the contribution of each cluster within the fear network and to disentangle the pattern of correlations between those and the activity of the lDLPFC, we observed the interaction graphs (Figure S2 in supplementary materials). Results showed that during the processing of the CS+ in late extinction, participants in the tDCS group presented a cross-cutting pattern of positive couplings with stronger correlations between the neural activity of the lDLPFC and the neural activity of the regions composing Cluster 1, when compared with the control group. Such stronger correlations were found between the lDLPFC and 64 AAL regions, including the left amygdala, the bilateral hippocampus, the bilateral insula, the bilateral caudate, the bilateral putamen, the bilateral pallidum, the bilateral thalamus, the bilateral cingulate cortex (anterior, middle and posterior), several regions of the inferior frontal cortex (left frontal triangular and orbital, left and right frontal medial orbital), and regions of the superior frontal cortex (including the right and left medial, the right and left frontal middle and the middle orbital frontal cortex), and the precuneus.

Cluster 2 included the left middle and left superior pole of the temporal lobe. Again, a pattern of stronger correlations after tDCS stimulation was found between these regions and the lDLPFC during the processing of the CS+ in late extinction.

The same was true for Cluster 3, which included the frontal middle, superior and inferior triangular parts of the left frontal cortex. Detailed results across the three clusters are depicted in Table S3 and Figure S2, in supplementary materials.

Besides the lDLPFC, no other phase or condition-dependent two-sample *t-tests* were statistically significant across other seeds.

### 3.4 Correlational analysis

To further clarify the association between day 1 and day 2 measures of fear response, we conducted correlational analysis between SCRs and self-report measures of arousal, valence, contingency and expectancy on day 1 and the neural activity on day 2.

We have found several correlations for the control group. Namely the neural activity of the left frontal superior region during the processing of the CS+ in late extinction, was positively correlated with day 1 self-reported contingency for the CS+ (r = .488, p = .047), with the SCRs during the processing of the CS+ and the CS-in the middle phase of the fear acquisition procedure (r = .671, p = .003 and r = .487, p = .041, respectively) and with the SCRs during the processing of the CS-in the late phase of the fear acquisition (r = .591, p = .012). Additionally, the SCRs differential (CS+ - CS-) in the middle phase of acquisition was positively correlated with the activity in the left frontal superior region during the processing of the CS+ in late extinction (r = .591, p = .012).

Furthermore, day 1 self-reported arousal for the CS+ was positively correlated with the percent signal change of the left frontal superior region during the processing of the CS- in late extinction (r = .649, p = .005) and negatively correlated with the percent signal change of the right paracentral region during the processing of the CS+ in late extinction (r = −.575, p = .016).

For the tDCS group, only pre-conditioning valence scores attributed to the CSs were found to be correlated with the activity of the right paracentral region during the processing of the CS+ (r = −.498, p = .049) and the processing of the CS- (r = .557, p = .025) both in late extinction.

## 4. Discussion

Understanding how conditioned fear is extinguished in experimental situations that mimic the therapeutic changes may contribute to the development of targeted clinical strategies for anxiety disorders. In our study, we examined how tDCS modulates neural responses and the connectivity within the fear pathway during a delayed extinction procedure. Due to known gender differences in what concerns anxiety responses (Boucsein W, Fowles DC, Grimnes S, Ben-Shakhar G, roth WT, Dawson ME, 2012) we tested only women who enrolled in two experimental sessions on two consecutive days (day 1 in the laboratory room, and day 2 inside the fMRI scanner).

On day 1, the SCRs as well as self-ratings confirmed that the fear conditioning procedure was weak but effective in building an associative fear memory and consequently in producing a conditioned fear response. As our hypothesis did not demand measures of successful extinction, in day 2 we did not collect other data than fMRI to measure how off-line tDCS interfered with the neural activity of the fear pathway.

We applied the cathode over the rDLPFC and the anode over the contralateral deltoid. We chose rDLPFC-contralateral deltoid montage, informed by previous preclinical studies that showed tDCS effectiveness in reducing anxiety related symptoms (Pedro Shiozawa et al., 2014a, 2014b). Hence, the DLPFC is known to be a hub region engaged during the processing of appraisal of fear experiences (Ertl et al., 2013), and the processing of fear extinction learning (Fullana et al., 2018). Furthermore, the rDLPFC is interconnected with the vmPFC to downregulate the amygdala during emotion regulation and value detection (Phelps et al., 2004).

Finally, as current modelling literature shows that increased current intensity does not occur under the electrode but rather between electrodes (Woods et al., 2016), we considered that the chosen tDCS montage could reliably answer our hypothesis according to which tDCS over the DLPFC allows for modulating the pathway that relies between electrodes and beyond. That is, we would be able to interfere not only with the cortical structures (such as the DLPFC, and other frontal, temporal, motor and parietal areas) but also with subcortical structures and the cerebellum.

Importantly, our fMRI contrast analysis results confirmed the study hypothesis, showing that tDCS over the rDLPFC interfered with delayed extinction learning by impacting the neural activity of two clusters during the processing of the CSs: Cluster 1 encompassed the frontal middle and the frontal superior left regions, and Cluster 2 encompassed the paracentral and postcentral areas.

In the tDCS group, Cluster 1 frontal regions showed a steeped decrease during the processing of the CS+ compared to controls (supporting a successful extinction learning about CS+ value), but an increased activity during the processing of the CS- which may be the neural correlate of the increased stimulus discrimination reported in previous studies (Dittert et al., 2018; Ganho-Ávila et al., 2019a). The increased activity for the CS-processing in late extinction though, additionally supports processes of reappraisal and detection value associated with uncertainty or with resistant extinction processes and which are typical of partial reinforcement schedules (Li et al., 2016; Rescorla, 1999). That is, whereas on day 1, the CS+ was learned to signal threat and the CS-was learned to be the safe cue, on day 2, the absence of information about the CS-(due to no US presentations) induced a state of uncertainty towards the CS-(for further discussion on processing of uncertainty, we recommend the recent work by Morriss and colleagues (Morriss et al., 2019)).

In Cluster 2 regions, the tDCS group showed an inverted pattern when compared to controls. For the tDCS group, the central and paracentral regions showed negative signal percentage changes during the processing of the CS+ (supporting a stronger extinction of the conditioned response), while simultaneously showing an increased activity during the processing of the CS-, similar to Cluster 1 pattern. This increased activation of the right central and post-central (somatosensory cortex [SI]) areas during the CS-is consistently reported during extinction learning studies (Fullana et al., 2018), supporting the hypothesis that tDCS enhances fear extinction-related processing. In fact, the SI is involved in the fear network through direct and indirect projections to the amygdala via insula for salient stimuli, participating in the generation of the emotion of fear, in processing of emotion significance, and in emotion regulation (Kropf et al., 2019).

Moreover, the findings of tDCS interference were further supported by correlational analysis that showed significant correlations between day 1 and day 2 measures and which were stimulation dependent. For example, for tDCS participants, those scoring the neutral stimuli more negatively in pre-acquisition showed an increased percent signal change in the right paracentral region on day 2 during the processing of the same neutral stimuli. Also, an increased post acquisition contingency for the CS+ correlated with an increased activity of the left frontal areas during the processing of the CS+ during extinction. These correlations were again stimulation-dependent, leading to the conclusion that tDCS is able to modulate the fear pathways and interrupt individual longitudinal patterns of fear response.

Interestingly, there were no differences found between groups with respect to the activity of core regions, such as the amygdalae and the hippocampi. This finding may be explained by one of two arguments: either the neural activity in both groups is equivalent in those regions and thus is stimulation-independent, or the lack of activity is due to methodological decisions that impact signal processing detection. In fact, an absent signal in the amygdala is frequently found due to the typically fast information processing of the ventral stream structures which may not be compatible with the parameters adopted to model its neural response. In fact, distinctive methodological options benefit the detection of neural signal in distinctive regions. For example, while modelling for shorter durations will likely capture subcortical processing, modelling for longer durations will lead to its loss (and the lack of differences detected in those regions) while offering information concerning cortical processes, such as memory and emotion regulation (Huettel et al., n.d.). Such may have been the case of our analysis.

The impact of tDCS on the fear pathway was definitely confirmed in our study with the results from contrast PPI analysis. We observed that cathodal stimulation of the rDLPFC increases the functional coupling between the lDLPFC and three clusters, during threat-specific processing in late extinction. This pattern of increased coupling and activation in the contralateral regions to the stimulation site has been previously reported in the literature. For example, Hanaoka and colleagues (2007) found a significant decrease in oxygenated haemoglobin during rTMS, followed by a significant increase in lDLPFC after low frequency rTMS (LF-rTMS) stimulation over the rDLPFC (Hanaoka et al., 2007). Both LF-rTMS and tDCS using cathodal stimulation over the rDLPFC inhibit neural excitability, and tDCS effects can last up to one hour (Stagg and Nitsche, 2011), which explains the contralateral effect we have found during late extinction.

Cluster 1 was the broader cluster detected in between group analysis, and included several AAL regions, all of which participating in the prefrontal cortical-amygdalo-hippocampal-cerebellar pathway: the hippocampi, the left amygdalae, the insulae, the basal ganglia, the thalamus and the anterior to posterior cingulate cortex, along with the inferior, middle and superior frontal cortex, up to the precuneus and finally reaching the anterior and medial regions of the cerebellum. This stronger coupling found in the tDCS group between those regions and the lDLPFC, during the processing of the CS+, suggests that tDCS effectively impacts the fear pathway synchrony. While it does not unequivocally inform in terms of direction of the information flow, it provides important clues. On the one hand, the synchrony of neural activity across the fear pathway during late extinction supports the neural correlate of an absence of inhibitory control towards subcortical structures, according to a topdown mechanism typical of classical extinction. On the contrary, such synchrony suggests memory updating through reconsolidation mechanisms which implicates increased plasticity in subcortical regions during the processing of threat. However, it would have given place to decreased fear responses after extinction, which we can not rule out. On the other hand, such lack of inhibitory control may explain the preservative long-term fear response towards the CS+, which we reported in our previous study (Ganho-Ávila et al., 2019a).

Also, because no group differences were found with respect to the processing of the CS-, we may argue that tDCS using cathodal electrode over the rDLPFC does not produce the generalization effect that has been described in the literature after stimulation with the anode electrode (Dittert et al., 2018), thus supporting the clinical benefits described in case reports (P. Shiozawa et al., 2014).

PPI contrast analysis also showed stronger time-course correlations between the lDLPFC and Cluster 2 regions during the processing of the CS+ in late extinction. Structures in the left temporal cortices such as the entorhinal cortex are crucial for memory consolidation of fear extinction (Izquierdo et al., 2006). As temporal regions serve as interface nodules between the amygdala, the hippocampus and the prefrontal cortex, our results show the increased coordination between frontal and temporal structures and support the use of delayed extinction when modulating consolidated fear memories.

The same stronger coupling was found between the lDLPFC and Cluster 3 during the processing of the CS+ in late extinction. This cluster included the left middle, the left superior and the left inferior triangular parts of the frontal cortex known to be involved in fear extinction and recall. In particular, previous literature shows the inferior frontal gyrus involvement in response inhibition (Hampshire et al., 2010), conflict resolution (Novick et al., 2005) and ambiguity (Lissek et al., 2020) which are cognitive functions recruited during the processing of the CS+ in late extinction allowing an efficient classical extinction process.

Here we show the online pattern of activity and connectivity for the prefrontal-cortical-subcortical-cerebellar pathway during the extinction processing of a previously consolidated fear memory. Additionally, we show that the relationship across the prefrontal-cortical-subcortical-cerebellar pathway is not inhibitory or anti-coupled in nature.

However, a number of limitations should be noted. Study results must not be generalized, as the sample involved only women given the gender differences across anxiety measures and fear conditioning responses (Lebron-Milad and Milad, 2012). Future replication studies should consider homogeneous samples of only men or heterogeneous samples to inform the field on whether and how gender interferes with the fear pathways. Additionally, in our study we did not control for the menstrual cycle phase and for the use of hormonal birth control methods. However, previous literature has showed menstrual cycle to impact the results concerning fear conditioning and extinction (Milad et al., 2010) and thus adequately considering controlling for these variables is deemed necessary in the future.

Second, we used a control group instead of a sham group, not blinding participants to the stimulation and not controlling for potential placebo effects. The use of a control group was intentional as a strategy to allow for increasing the recruitment rate by including participants not willing to do tDCS, and participants not complying with tDCS safety criteria. In fact, although this was not the first option for the study design, it was a much-needed decision after the authors realized the significative number of participants that were willing to be part of the study but not comfortable to complete tDCS sessions. Furthermore, a recent tDCS-based experimental study (Rabipour et al., 2018) showed that the expectations primed to participants concerning stimulation outcomes account for most of the tDCS effects. In our study, participants were naïve to the procedures and naïve to the study hypothesis, bearing it difficult for any personal expectations to have a significative impact in results.

Third, the acquisition effect as indexed by SCRs was weak which may be claimed as hardly supporting a successful fear acquisition and subsequent extinction procedure. However, SCRs were not the only measures collected to show fear acquisition in day 1, and self-report measures confirm that the learning of fear acquisition was achieved. Furthermore, our correlational data between day 1 and day 2 measures exclusively for the control group showed the dependency of day 2 neuronal activity during extinction in relation to day 1 self-reports and SCRs, further supporting successful fear acquisition.

Forth, although our hypothesis did not demand measures of successful extinction, we cannot state that the patterns of neural activity observed directly matched successful extinction learning. Nonetheless, our focus is on the ongoing delayed extinction and we aim to advance the field by offering the information about the impact of tDCS using the cathode electrode over the rDLPFC and its use as a complementary strategy to exposure-based treatments. In future studies, however, simultaneous measures of extinction (e.g. SCRs and self-reports of contingency and expectancy) should be collected inside the scanner to address lack of longitudinal data.

Fifth, changing contexts from day 1 (laboratory) to day 2 (MRI scanner), can be a limitation as context is known to modulate fear extinction responses (Bouton, 2002, 1993). Future studies should conduct all sessions in the same context or control for the change of context with additional experimental arms.

### 4.1 Conclusions

To date, most studies in fear extinction have failed to appreciate the importance of memory consolidation during the extinction-learning phase. Our results offer a first understanding of the effect of tDCS with the cathode electrode over the rDLPFC and extracephalic anode in extinction, supporting its clinical use. Here, we compared the mechanisms at play during extinction *vs.* extinction combined with tDCS where stimulation was used to inhibit the rDLPFC, a crucial hub to emotion regulation processes. Our data suggests that tDCS increased the processing of uncertainty associated with the safety cue, by modulating frontal regions. Additionally, after tDCS the time course activity correlation between regions of the fear pathway was stronger during the processing of threat.

More importantly, as this experimental setting mimics real-life context changes, by allowing consolidation mechanisms to take place, we can state that targeting this regulatory system is a promising venue to non-invasive brain stimulation approaches for the treatment of anxiety disorders.

The question of whether the tDCS combined with exposure-based strategies is more successful via the stimulation of the DLPFC using the cathode electrode or the anode electrode should be targeted in future studies.

## DECLARATIONS

### Funding

This work was supported by a Foundation for Science and Technology of Portugal and Programa COMPETE grant to AGA and JA [grant numbers PTDC/MHC-PCN/3575/2012, SFRH/BD/80945/2011, PTDC/MHC-PAP/5618/2014 (POCI-01-0145-FEDER-016836)]. The Psychology Research Centre of Minho University is supported by the Portuguese Foundation for Science and Technology and the Portuguese Ministry of Education and Science through national funds and co-financed by FEDER through COMPETE2020 under the PT2020 Partnership Agreement [POCI-01-0145-FEDER-007653].

The Cognitive and Behavioural Centre for Research and Intervention

Of the Faculty of Psychology and Educational Sciences of the University of Coimbra is supported by the Portuguese Foundation for Science and Technology and the

Portuguese Ministry of Education and Science through national funds and co-financed by

FEDER through COMPETE2020 under the PT2020 Partnership

Agreement [UID/PSI/01662/2013].

## Ethics approval

The current study was approved by the ethical committee of the Faculty of Psychology and

Educational Sciences of the University of Coimbra

## Consent to participate (include appropriate statements)

Participants gaven their informed consent to participate according to the local ethical committee recommendations (compliant with the Declaration of Helsinki and GDPR recommendations).

## Availability of data and material

In line with Open Science practices, fMRI, SCR and Self reports raw data, scripts for stimuli presentation, preprocessing and analysis may be found at https://osf.io/umy7c/

## Code availability

Not applicable

## Authors’ contributions

**Ana Ganho-Ávila:** Conceptualization, methodology, investigation, software, formal analysis, visualization, supervision, verification, funding acquisition, project administration, writing- original draft preparation

**Raquel Guiomar**: Investigation, software, formal analysis, data curation, writing- reviewing and editing,

**Daniela Valério**: Software, formal analysis, writing- reviewing and editing,

**Óscar Gonçalves:** Conceptualization, methodology, supervision, funding acquisition, writing-revie wing and editing,

**Jorge Almeida** : Conceptualization, methodology, software, supervision, funding acquisition, writing- revie wing and editing.

## Acknowledgements

We thank Soares, M.J, Gerardo, B. for data collection and pre-processing. We Thank Professor Karl Friston for his wise advice in methodological choices and analysis.

## Sample CRediT author statement

**Ana Ganho-Ávila:** Conceptualization, methodology, investigation, software, formal analysis, visualization, supervision, verification, funding acquisition, project administration, writing-original draft preparation

**Raquel Guiomar**: Investigation, software, formal analysis, data curation, writing-reviewing and editing,

**Daniela Valério**: Software, formal analysis, writing-reviewing and editing,

**Óscar Gonçalves:** Conceptualization, methodology, supervision, funding acquisition, writing-reviewing and editing,

**Jorge Almeida**: Conceptualization, methodology, software, supervision, funding acquisition, writing-reviewing and editing,

