## Supplementary materials for "Offline tDCS modulates prefrontal-cortical-subcortical-cerebellar fear pathways in delayed fear extinction"

^a^Proaction Laboratory, Faculty of Psychology and Educational Sciences, University of Coimbra, 3001-802 Coimbra, Portugal; ^b^Center for Research in Neuropsychology and Cognitive Behavioral Intervention – CINEICC of the Faculty of Psychology and Educational Sciences, University of Coimbra, 3001-802 Coimbra, Portugal _;_^c^Clinical & Developmental Neurosciences, CIPSI, University of Minho, Braga; ^d^School of Psychology, University of Minho, 4710-057 Braga, Portugal; ^e^Department of Applied Psychology, Northeastern University, Boston, Massachusetts, USA.

**^*^Corresponding author:** Ana Ganho-Ávila

Address: ^b^Center for Research in Neuropsychology and Cognitive Behavioral Intervention – CINEICC, Faculty of Psychology and Educational Sciences, University of Coimbra, Rua do Colégio Novo, 3000-115 Coimbra, Portugal

This document includes:

- Tables: S1, S2, S3, S4.

- Figures: S1, S2.

In line with Open Science practices, fMRI, SCR and Self reports raw data, scripts for stimuli presentation, preprocessing and analysis may be found at [https://osf.io/umy7c/](https://www.google.com/url?q=https://osf.io/umy7c/&sa=D&source=hangouts&ust=1576752421796000&usg=AFQjCNFRIBesynovQM57-UIPWUOybGHswg) (Doi: 10.17605/OSF.IO/UMY7C).

| **Table S1**. *Participants Sociodemographic and clinical characteristics.* | | | | | | | |
| --- | --- | --- | --- | --- | --- | --- | --- |
|  | **N** | **Cathodal group** | **N** | **Control group** | | ***p*** | ***df*** |
| **Age, Years** | 16 | 24.19 (5.94) | 18 | 22.56 (5.47) | .410 | | 32 |
| **Education, Years** | 16 | 15.25 (2.96) | 18 | 15.00 (3.14) | .813 | | 32 |
| **US Intensity, %** | 16 | 90.80 (3.13) | 16 | 93.44 (4.50) | .061 | | 31 |
| **ASI-3-PT** | 16 | 19.13 (12.10) | 15 | 21.40 (14.41) | .519 | | 31 |
| **BSI (GSI)** | 15 | .028 (.007) | 14 | .028 (.006) | .967 | | 27 |
| **STAI 1** | 16 | 37.13 (13.08) | 16 | 35.56(10.75) | .925 | | 32 |
| **STAI 2** | 16 | 40.06 (11.94) | 16 | 38.44 (11.37) | .861 | | 32 |
| Note. Mean values are depicted. The standard deviations (SD) are within parentheses. US: Unconditioned stimuli; ASI-3-PT: Anxiety Sensitivity Scale Portuguese version; BSI (GSI): Global index for symptoms intensity of the Behavioural Symptoms Inventory; STAI 1: State Anxiety Inventory; STAI 2: Trace Anxiety Inventory. | | | | | | | |

**Table S2.** *tDCS side effects.*

| No side effects | 37,50 % |
| --- | --- |
| Itching arm (skin under anodal electrode) | 37,50 % |
| Tingling (skin under anodal electrode) | 18,75 % |
| Sleepiness | 18,75 % |
| Redness of the skin (skin under cathodal electrode) | 12,50 % |
| Headache | 6,25 % |

**Table S3.** *Whole brain contrast analysis.*

| Peak MNI coordinate region | Peak voxel  MNI coordinates | | | Number of voxels | Peak Z | Mean Z |
| --- | --- | --- | --- | --- | --- | --- |
|  | **X** | **Y** | **Z** |  |  |  |
| Contrasts two sample t test, CS+>CS- late phase | | | | | | |
| Cluster 1 – Frontal Sup L | -18 | 15 | 45 | 85 | 2.18 | 2.07 |
| Cluster 2 – Paracentral Lobule R | 9 | -33 | 57 | 43 | 2.02 | 2.00 |

Note: Group level significance established at p < 0.05 (TFCE corrected at the voxel level; clusters did not survive Bonferroni corrections for multiple comparisons).

**Table S4.** *PPI analysis, clusters and subdivisions.*

| Peak MNI coordinate region | Peak voxel  MNI coordinates | | | Number of voxels | Peak Z | Mean Z |
| --- | --- | --- | --- | --- | --- | --- |
|  | **X** | **Y** | **Z** |  |  |  |
| PPI two sample t test, seed: lDLPFC to grey matter mask, CS+>CS- late phase | | | | | | |
| Cluster 1 – Frontal Sup R | **24** | **54** | **9** | **7055** | **-3.54** | **-2.89** |
| Amygdala_L | -28 | -6 | -12 | 9 | -3.22 |  |
| Calcarine_L | 2 | -64 | 14 | 28 | -3.09 |  |
| Calcarine_R | 12 | -60 | 10 | 124 | -3.16 |  |
| Caudate_L | -16 | 12 | 8 | 92 | -3.40 |  |
| Caudate_R | 18 | 16 | 4 | 112 | -3.35 |  |
| Cerebelum_3_L | -6 | -46 | -20 | 11 | -3.08 |  |
| Cerebelum_3_R | 12 | -44 | -24 | 11 | -2.94 |  |
| Cerebelum_4_5_L | -10 | -50 | -20 | 85 | -3.09 |  |
| Cerebelum_4_5_R | 10 | -54 | -4 | 58 | -3.03 |  |
| Cerebelum_6_L | -24 | -50 | -26 | 37 | -2.94 |  |
| Cerebelum_6_R | 18 | -56 | -28 | 18 | -2.90 |  |
| Cerebelum_9_R | 12 | -45 | -36 | 15 | -2.94 |  |
| Cingulum_Ant_L | -4 | 24 | 30 | 288 | -3.35 |  |
| Cingulum_Ant_R | 12 | 40 | 16 | 288 | -3.35 |  |
| Cingulum_Mid_L | -4 | 24 | 32 | 139 | -3.35 |  |
| Cingulum_Mid_R | 4 | -18 | 34 | 251 | -3.42 |  |
| Cingulum_Post_L | -10 | -48 | 18 | 46 | -2.97 |  |
| Cingulum_Post_R | 6 | -42 | 18 | 42 | -3.16 |  |
| Cuneus_R | 20 | -62 | 20 | 7 | -3.05 |  |
| Frontal_Inf_Oper_R | 42 | 18 | 30 | 114 | -3.43 |  |
| Frontal_Inf_Orb_L | -34 | 18 | -14 | 166 | -3.36 |  |
| Frontal_Inf_Orb_R | 36 | 24 | -18 | 140 | -3.35 |  |
| Frontal_Inf_Tri_L | -36 | 24 | -2 | 61 | -3.35 |  |
| Frontal_Inf_Tri_R | 42 | 18 | 28 | 328 | -3.43 |  |
| Frontal_Med_Orb_L | -12 | 42 | -8 | 12 | -2.97 |  |
| Frontal_Med_Orb_R | 12 | 52 | -2 | 64 | -3.34 |  |
| Frontal_Mid_L | -20 | 46 | 30 | 110 | -3.24 |  |
| Frontal_Mid_R | 26 | 54 | 20 | 452 | -3.47 |  |
| Frontal_Mid_Orb_R | 30 | 48 | -4 | 11 | -2.63 |  |
| Frontal_Sup_Medial_L | -10 | 24 | 36 | 207 | -3.35 |  |
| Frontal_Sup_Medial_R | 12 | 52 | 0 | 220 | -3.35 |  |
| Frontal_Sup_L | -12 | 26 | 38 | 88 | -3.33 |  |
| Frontal_Sup_R | 24 | 54 | 10 | 328 | -3.54 |  |
| Fusiform_L | -26 | -48 | -10 | 19 | -3.06 |  |
| Fusiform_R | -26 | -48 | -10 | 17 | -3.06 |  |
| Heschl_L | -36 | -30 | 12 | 25 | -2.95 |  |
| Heschl_R | 42 | -26 | 6 | 28 | -2.77 |  |
| Hippocampus_L | -24 | -30 | -6 | 33 | -3.16 |  |
| Hippocampus_R | 20 | -36 | 0 | 56 | -3.24 |  |
| Insula_L | -32 | 12 | -8 | 249 | -3.43 |  |
| Insula_R | 34 | 20 | -14 | 144 | -3.34 |  |
| Lingual_L | -24 | -50 | -8 | 86 | -3.09 |  |
| Lingual_R | 12 | -54 | 0 | 98 | -3.16 |  |
| Pallidum_L | -18 | 6 | 4 | 32 | -3.37 |  |
| Pallidum_R | 18 | 8 | 4 | 33 | -3.21 |  |
| Paracentral_Lobule_R | 6 | -34 | 52 | 11 | -2.87 |  |
| ParaHippocampal_L | -22 | -36 | -6 | 26 | -3.03 |  |
| ParaHippocampal_R | 22 | -36 | -6 | 76 | -3.18 |  |
| Postcentral_L | -48 | -6 | 18 | 6 | -2.33 |  |
| Precuneus_L | -18 | -50 | 0 | 34 | -3.06 |  |
| Precuneus_R | 10 | -54 | 8 | 136 | -3.13 |  |
| Putamen_L | -28 | 12 | -6 | 201 | -3.43 |  |
| Putamen_R | 24 | 12 | 6 | 166 | -3.35 |  |
| Rolandic_Oper_L | -42 | -6 | 18 | 26 | -2.39 |  |
| Supp_Motor_Area_R | 8 | 20 | 48 | 25 | -3.01 |  |
| Temporal_Mid_L | -62 | -14 | -6 | 26 | -2.53 |  |
| Temporal_Sup_L | -42 | -30 | 6 | 66 | -2.95 |  |
| Temporal_Sup_R | 50 | -22 | 4 | 43 | -2.78 |  |
| Thalamus_L | -16 | 12 | 8 | 161 | -3.40 |  |
| Thalamus_R | 12 | -14 | 0 | 147 | -3.29 |  |
| Vermis_1_2 | 0 | -40 | -20 | 11 | -2.93 |  |
| Vermis_3 | 0 | -48 | -18 | 15 | -3.09 |  |
| Vermis_4_5 | 6 | -54 | 2 | 42 | -3.14 |  |
| Cluster 2 – Temporal Mid L | **-44** | **4** | **-24** | **38** | **-2.37** | **-2.34** |
| Temporal_Mid_L | -44 | 4 | -24 | 19 | -2.37 |  |
| Temporal_Pole_Sup_L | -44 | 6 | -24 | 12 | -2.36 |  |
| Cluster 3 – Frontal Mid L | **-30** | **30** | **36** | **88** | **-2.40** | **-2.29** |
| Frontal_Mid_L | -30 | 30 | 36 | 68 | -2.40 |  |
| Frontal_Inf_Tri_L | -36 | 18 | 26 | 12 | -2.31 |  |
| Frontal_Sup_L | -28 | 32 | 34 | 5 | -2.34 |  |

Note: *Regions showing statistically significant differences between groups are depicted (TFCE corrected at the voxel level and Bonferroni corrected accounting for the number of contrasts). Clusters and subdivisions as per AAL regions.*

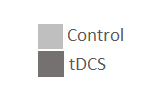

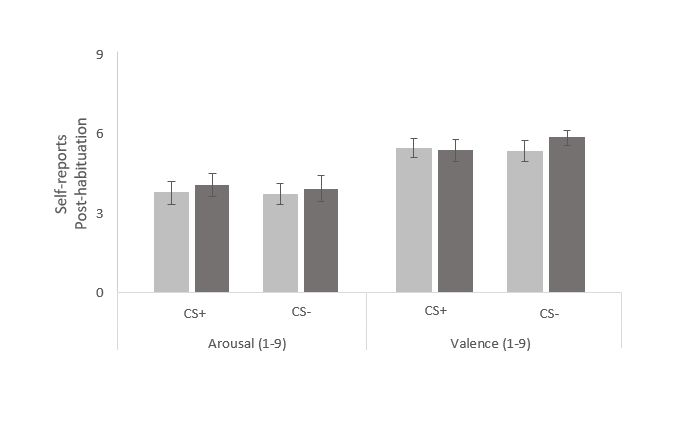

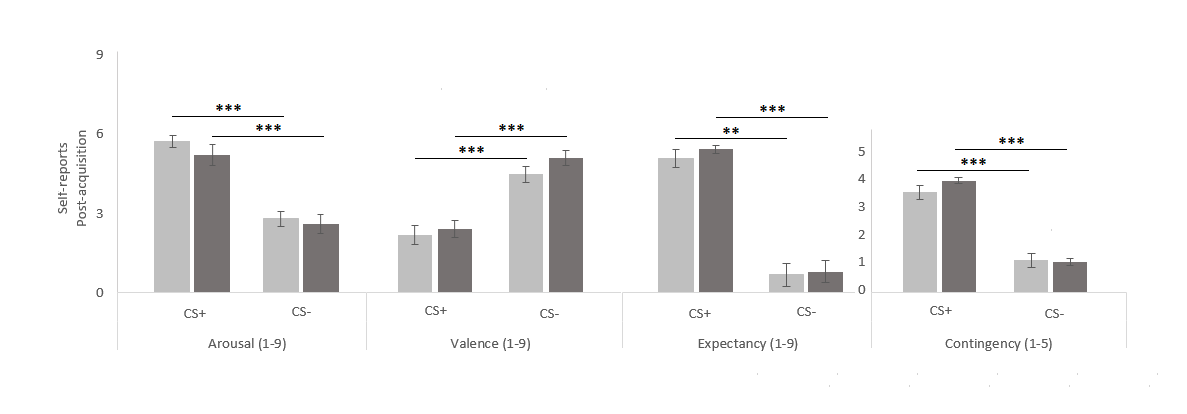

**Figure S1. Day 1 Self-reports.** Post-Habituation Self-reports of arousal and valence (top). Post-Acquisition arousal, valence, expectancy and contingency (bottom). CS+: conditioned stimuli; CS-: non-reinforced or control stimuli; tDCS: tDCS cathodal stimulation group; Control: control group. Error bars represent standard errors of the mean (SEM). **p < .01, ***p < .001.

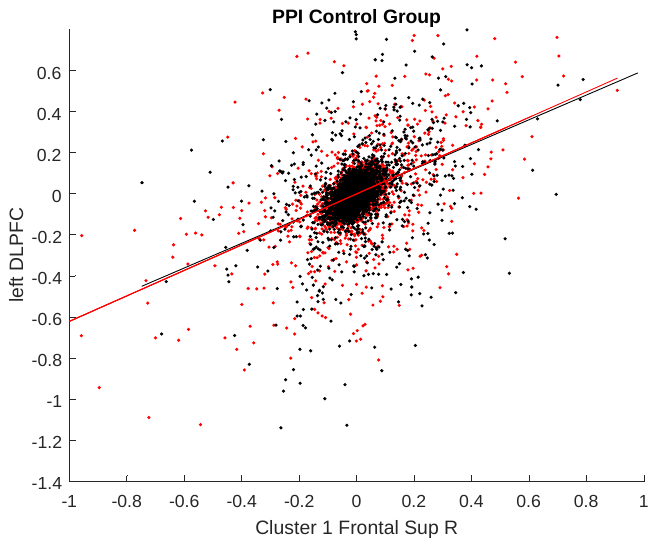

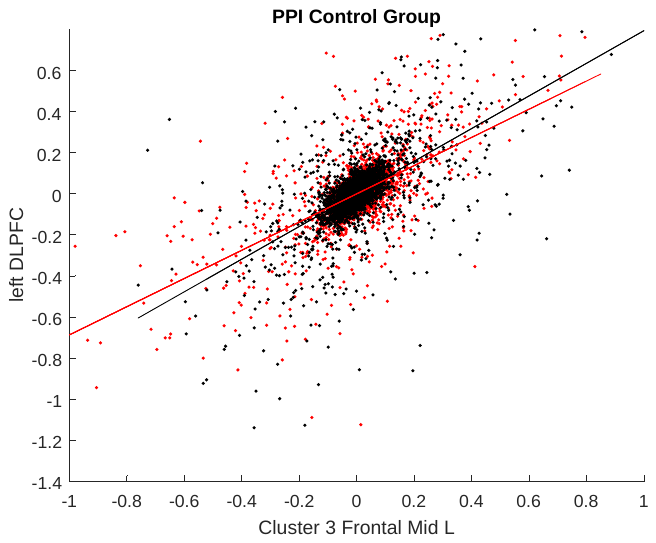

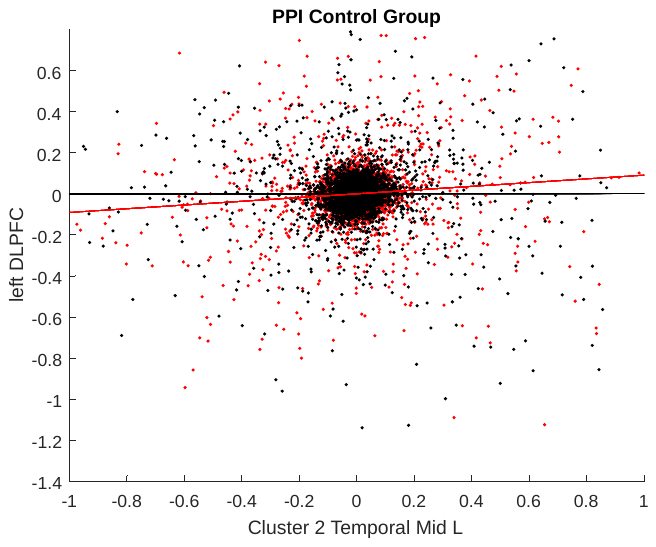

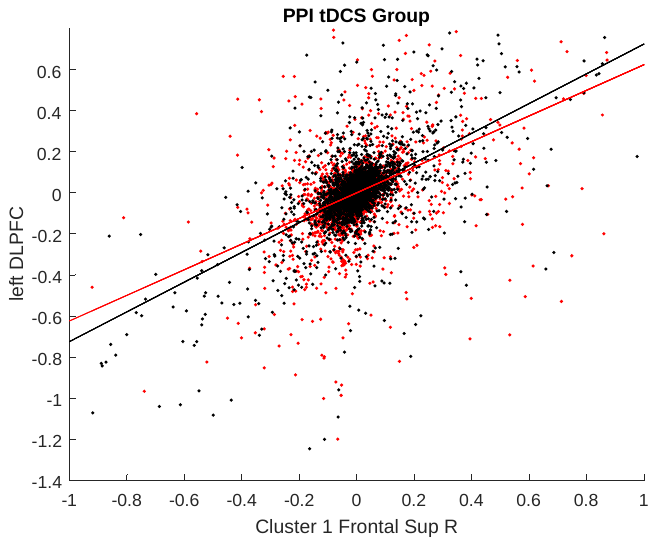

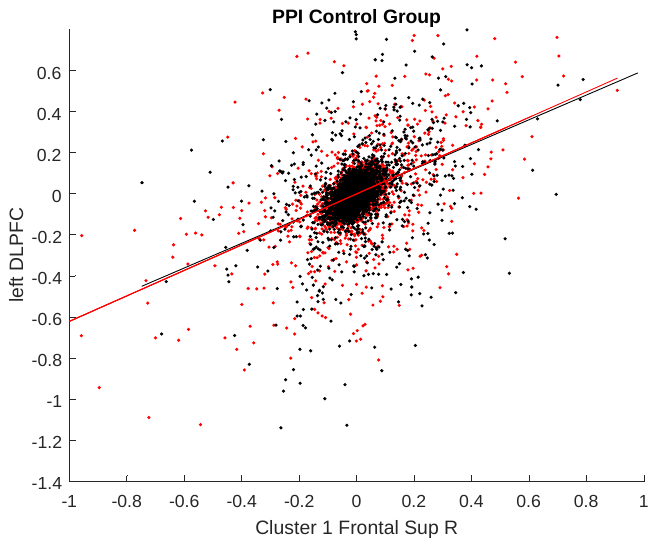

| 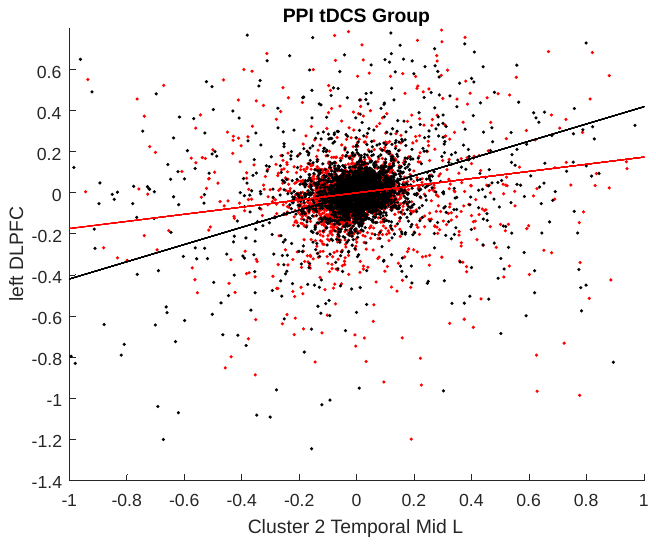 | 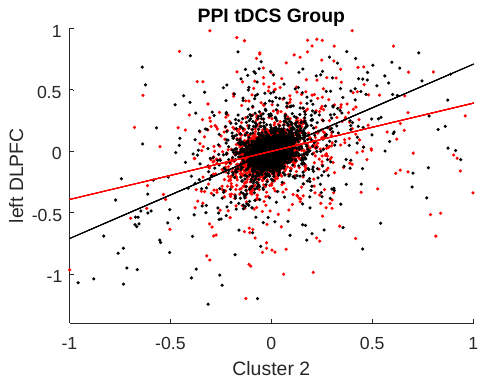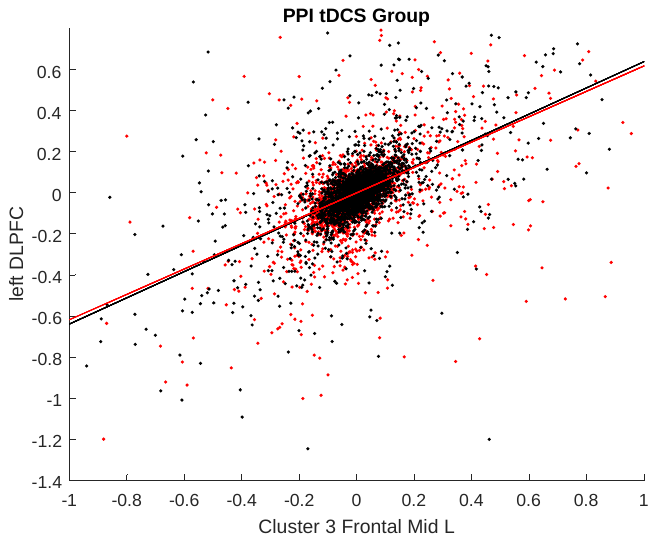 | 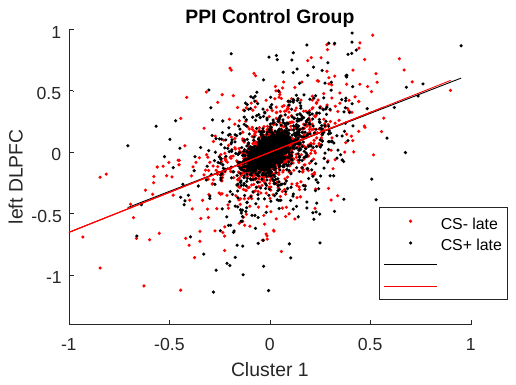 |
| --- | --- | --- |

**Figure S2. Graphical representation PPI two-sample t test for clusters 1, 2, and 3.** Group level significance was established at p < 0.05 (TFCE corrected; thresholded at z > 2.24*)* and between group comparisons were Bonferroni corrected accounting for four contrasts*.* Seed was the left DLPFC, reference voxels were whole brain grey matter mask, behavioral contrast was CS+>CS- in the late phase of the fear extinction session (second half of the task).
